# The dilution effect limits plasmid horizontal transmission in multispecies bacterial communities

**DOI:** 10.1101/2021.05.18.444624

**Authors:** Anastasia Kottara, Laura Carrilero, Ellie Harrison, James P.J. Hall, Michael A. Brockhurst

**Author notes:** Corresponding author: Michael Brockhurst, Division of Evolution and Genomic Sciences, Faculty of Biology, Medicine and Health, University of Manchester, Manchester, M13 9PT, UK.

## Abstract

By transferring ecologically important traits between species, plasmids drive genomic divergence and evolutionary innovation in their bacterial hosts. Bacterial communities are often diverse and contain multiple coexisting plasmids, but the dynamics of plasmids in multispecies communities are poorly understood. Here, we show, using experimental multispecies communities containing two plasmids, that bacterial diversity limits the horizontal transmission of plasmids due to ‘the dilution effect’; an epidemiological phenomenon whereby living alongside less proficient host species reduces the expected infection risk for a focal host species. In addition, plasmid horizontal transmission was also affected by plasmid diversity, such that the rate of plasmid conjugation was reduced from coinfected host cells carrying both plasmids. In diverse microbial communities, plasmid spread may be limited by the dilution effect and plasmid-plasmid interactions reducing the rate of horizontal transmission.

## Introduction

Mobile genetic elements (MGE) are an important source of potentially beneficial accessory traits for host bacteria, equipping these bacterial cells with new ready-to-use functions and thereby allowing them to expand their ecological niche [1–3]. Plasmids are common in bacterial communities, infecting diverse bacterial taxa [4], and often multiple plasmids co-exist in natural microbial communities [5–6]. The long-term persistence of plasmids in bacterial communities will depend both on the proficiency of host species to stably maintain plasmids in their populations by vertical transmission [7], and the rate of horizontal transmission of plasmids within and between species by conjugation [8].

Previous studies have shown that plasmids are not equally maintained across different host species [9–10], while plasmid transmission dynamics are affected by bacterial community structure [8]. Thus, in communities where plasmids rely on horizontal transmission for their maintenance [8,11], plasmid dynamics could be affected by the diversity of the community, especially if the different host species differ in their proficiency and transmission rates. Studies focused on parasite transmission in host communities have shown that the transmission of multi-host parasites can be limited by species richness, which is termed the ‘dilution effect’ [12–13]: A focal host species has a reduced risk of parasite infection when in a diverse community than would be expected from its intraspecific transmission rate, if transmission from other species in the community is less efficient [14]. We hypothesise that the dilution effect may also apply to plasmids in communities where hosts differ in their ability to maintain and transmit plasmids.

To gain a better understanding of plasmid dynamics in complex multi-plasmid / multi-host communities we constructed simple bacterial communities in effectively sterile potting soil (soil microcosms) under controlled laboratory conditions and tracked plasmid dynamics over-time. Specifically, communities contained two distinct conjugative plasmids, pQBR57 and pQBR103, that are known to vary in their rate of conjugation within populations of the focal species, *P. fluorescens* SBW25 [15]. *P. fluorescens* SBW25 populations were embedded within a community of five *Pseudomonas* species, and these were compared to controls where *P. fluorescens* SBW25 was propagated in monoculture. We report that presence of the *Pseudomonas* community reduced the rate of plasmid co-infection in *P. fluorescens* SBW25 in line with there being a dilution effect limiting the rate of horizontal transmission in more diverse communities.

## Materials and methods

### Bacterial strains and plasmids

*P. fluorescens* SBW25 [16] was the plasmid-donor in this study, carrying either the plasmid pQBR57 or pQBR103. *P. fluorescens* SBW25 was labelled by directed insertion of gentamicin resistance (Gm^R^) as previously described [17]. The plasmids used in this study, pQBR103 and pQBR57 are large conjugative plasmids (425 kb and 307 kb respectively) that confer mercury resistance via a *mer* operon encoded on a Tn5042 transposon [5, 15, 18]. Both plasmids were independently conjugated into gentamicin resistant (Gm^R^) *P. fluorescens* SBW25 from streptomycin resistant (Sm^R^) plasmid-bearing *fluorescens* SBW25. pQBR57 was also conjugated from *P. fluorescens* SBW25 Sm^R^ into *P. fluorescens* SBW25(pQBR103) Gm^R^ in order to obtain *P. fluorescens* SBW25 (pQBR103:pQBR57). Each plasmid-donor was mixed in 1:1 ratio with the plasmid-recipient strain, incubated for 48 h and spread on King’s B (KB) agar plates containing 10 μg ml^−1^ gentamicin and 20 μM of mercury(II) chloride to select for transconjugant colonies [19]. As previously described, the conjugation assays were conducted in 6 ml KB growth medium in 30 ml universal vials (‘microcosms’) at 28°C in shaking conditions (180 rpm). Background communities consisted of five different *Pseudomonas* species; *P. stutzeri* JM300 (DSM 10701) [20], *P. putida* KT2440 [21], *P. protegens* Pf-5 [22], *P. fluorescens* Pf0-1 [23], *P. aeruginosa* PAO1 [24].

### Selection experiment

Twelve colonies of the plasmid-bearing *P. fluorescens* SBW25(pQBR103) and *P. fluorescens* SBW25(pQBR57) were grown overnight in KB microcosms at 28°C with shaking 180 rpm. Six colonies of each of the plasmid-free *Pseudomonas* species (*P. stutzeri* JM300 (DSM 10701), *P. putida* KT2440, *P. protegens* Pf-5, *P. fluorescens* Pf0-1, *P. aeruginosa* PAO1) were also grown overnight in KB microcosms using the same culture conditions. Six replicate populations containing equal proportions of *P. fluorescens* SBW25(pQBR103) and *P. fluorescens* SBW25(pQBR57) were propagated either with or without the background community of five *Pseudomonas* species. Populations were grown in potting soil microcosms supplemented with mercury (16 μg g^−1^ Hg(II)). Each community had a starting ratio of 1:1 between *P. fluorescens* SBW25(pQBR103) and *P. fluorescens* SBW25(pQBR57) (~each 1×10^6^ cfu g^−1^) such that the starting frequencies of pQBR103 and pQBR57 were approximately 50%. The background community of *Pseudomonas* species contained each species in equal proportion (~each 4×10^5^ cfu g^−1^). To prepare the soil inoculum, the mix of each community (final volume: 100 μl) was centrifuged for 1 min at 10,000 rpm and resuspended in 1 ml M9 salt solution [25]. Next, the soil microcosms (10 g twice-autoclaved John Innes No. 2 compost soil) were inoculated with 100 μl of the mix, briefly vortexed to disperse the inoculum in the soil, and incubated at 28°C at 80% humidity [8]. Every 4 days, 10 ml of M9 buffer and 20 glass beads were added to each soil microcosm and mixed by vortexing for 1 min, and 100 μl of soil wash was transferred to a fresh soil microcosm as previously described by Hall *et al.* [8]. The communities were propagated for 6 transfers (24 days, estimated to be approx. 42 bacterial generations).

At each transfer, total population counts were estimated by plating onto non-selective KB agar plates. Bacterial counts for the plasmid-bearing *P. fluorescens* SBW25 strains were estimated by plating onto selective media: 10 μg ml^−1^ gentamicin KB agar plates. Each of these plates were then replica plated onto mercury KB agar plates (100 μM mercury(II) chloride) in order to assess the frequency of mercury resistance within *P. fluorescens* SBW25 and at the whole-community level. Twenty-four colonies of *P. fluorescens* SBW25 were sampled every 2 transfers from the mercury containing plates and tested for the presence of the plasmids and mercury transposon by PCR screening. Twenty-four colonies of the total community were randomly sampled from the mercury containing plates at two time-points (transfers 4 and 6) and also tested for the presence of the plasmids and mercury transposon. The PCR screening was designed to use three set of primers that targeted the *mer* operon-Tn5042 transposon [forward primer: 5′-TGCAAGACACCCCCTATTGGAC-3′, reverse primer: 5′-TTCGGCGACCAGCTTGATGAAC-3′], the pQBR103-plasmid specific origin of replication *oriV* [forward primer: 5′-TGCCTAATCGTGTGTAATGTC-3′, reverse primer: 5′-ACTCTGGCCTGCAAGTTTC-3′] and the pQBR57-plasmid specific *uvrD* gene [forward primer: 5′-CTTCGAAGCACACCTGATG-3′, reverse primer: 5′-TGAAGGTATTGGCTGAAAGG-3′] [26].

### Competitive fitness assay

Four individual colonies of the ancestral *P. fluorescens* SBW25(pQBR103:pQBR57) were competed against the plasmid-free *P. fluorescens* SBW25 with and without the five-species community. The fitness assay was performed with and without mercury in soil microcosms. Relative fitness was measured by mixing differentially the plasmid-bearer (Gm^R^) and plasmid-free (Sm^R^) in 1:1 ratio. The five-species community was added in the same ratio as at the beginning of the selection experiment. The inoculum was diluted 100-fold in M9 salts before being added into soil microcosms and incubated at 28°C and 80% humidity for 4 days. Samples were plated on KB agar plates supplemented with selective concentration of 10 μg ml^−1^ gentamicin and 50 μg ml^−1^ streptomycin at the beginning and end of the competition to estimate the density of plasmid-bearing and plasmid-free bacteria. The relative fitness was calculated as the selection rate (r) [27].

### Conjugation assay

Four individual colonies of each ancestral *P. fluorescens* SBW25 (pQBR103:pQBR57), *P. fluorescens* SBW25 (pQBR103) and *P. fluorescens* SBW25 (pQBR57) were conjugated into the isogenic plasmid-free strain. Conjugation rate of the different plasmids was measure by mixing differentially the plasmid-bearer (Gm^R^ or Sm^R^) and plasmid-free (Sm^R^ or Gm^R^ respectively) in 1:1 ratio. The mix was centrifuged for 1 min at 10,000 rpm to remove spent media, resuspended in M9 salt solution, diluted 100-fold in high (KB), medium (0.1x KB) and low (0.01x KB) resource media and incubated at 28°C for 48 h. KB agar plates were supplemented with 10 μg ml^−1^ gentamicin or 50 μg ml^−1^ streptomycin to estimate the density of plasmid-donor and plasmid-recipient bacteria at the beginning and end of the assay. KB agar plates were supplemented with 10 μg ml^−1^ gentamicin and 20 μM mercury(II) chloride or 50 μg ml^−1^ streptomycin and 20 μM mercury(II) chloride to estimate the density of the transconjugant bacteria at the end of the assay. The conjugation rate was calculated with the method firstly described by Simonsen *et al*. [19].

### Statistical analyses

Statistical analyses were performed using RStudio version 3.2.3 [28]. The prevalence of each plasmid status (pQBR103 only, pQBR57 only, or both) in *P. fluorescens* SBW25 was estimated as the area under the curve using the function *auc* of the package ‘flux’ [29]. One-way ANOVA tests compared plasmid prevalence in *P. fluorescens* SBW25 with versus without the community. Kruskal-Wallis test was used to analyse the end-point frequency of each plasmid at a whole-community level since the data was not normally distributed. Welch’s t-test was used to analyse the effect of the background community on the relative fitness of *P. fluorescens* SBW25 carrying both plasmids. Kruskal-Wallis test was used to assess the differences between the conjugation rates of pQBR57 and pQBR57: pQBR103 in the different resource media as the data were not normally distributed; the conjugation rate of pQBR103 plasmid was not in detectable range in medium and low resource media thus pQBR103 was not included in this statistical analysis. Welch’s t-test was used to compare the conjugation rate of pQBR57 to pQBR57:pQBR103 and pQBR103 plasmid in high resource media where the conjugation rate of each plasmid was in detectable range.

## Results

### Plasmid co-infection limited in community

While mercury resistance remained at fixation in all replicates, we observed contrasting plasmid dynamics in the *P. fluorescens* SBW25 population with versus without the background *Pseudomonas* community. In the presence of the background community, in the majority of replicates the *P. fluorescens* SBW25 population was dominated by pQBR103, such that bacteria were typically either singly-infected by pQBR103 or co-infected with both pQBR103 and pQBR57. By contrast, in the absence of the background community we observed higher rates of co-infection with both pQBR103 and pQBR57, or, in a single replicate, the fixation of pQBR57. Overall, we observed that the frequency of plasmid co-infection was higher in the absence of the background community (ANOVA F_1,10_=5.569, p=0.039; Figure 1). To test if this effect could be caused by higher fitness costs of plasmid co-infection in the presence versus absence of the community, perhaps due to more intense resource competition, we competed *P. fluorescens* SBW25(pQBR103:pQBR57) against plasmid-free *P. fluorescens* SBW25 with or without the background community. We found, however, that the presence of the background community had no effect on the relative fitness of *P. fluorescens* SBW25(pQBR103:pQBR57) (Welch’s t-test, t_13.68_=0.698, p=0.496; Figure 2).

**Figure 1.**
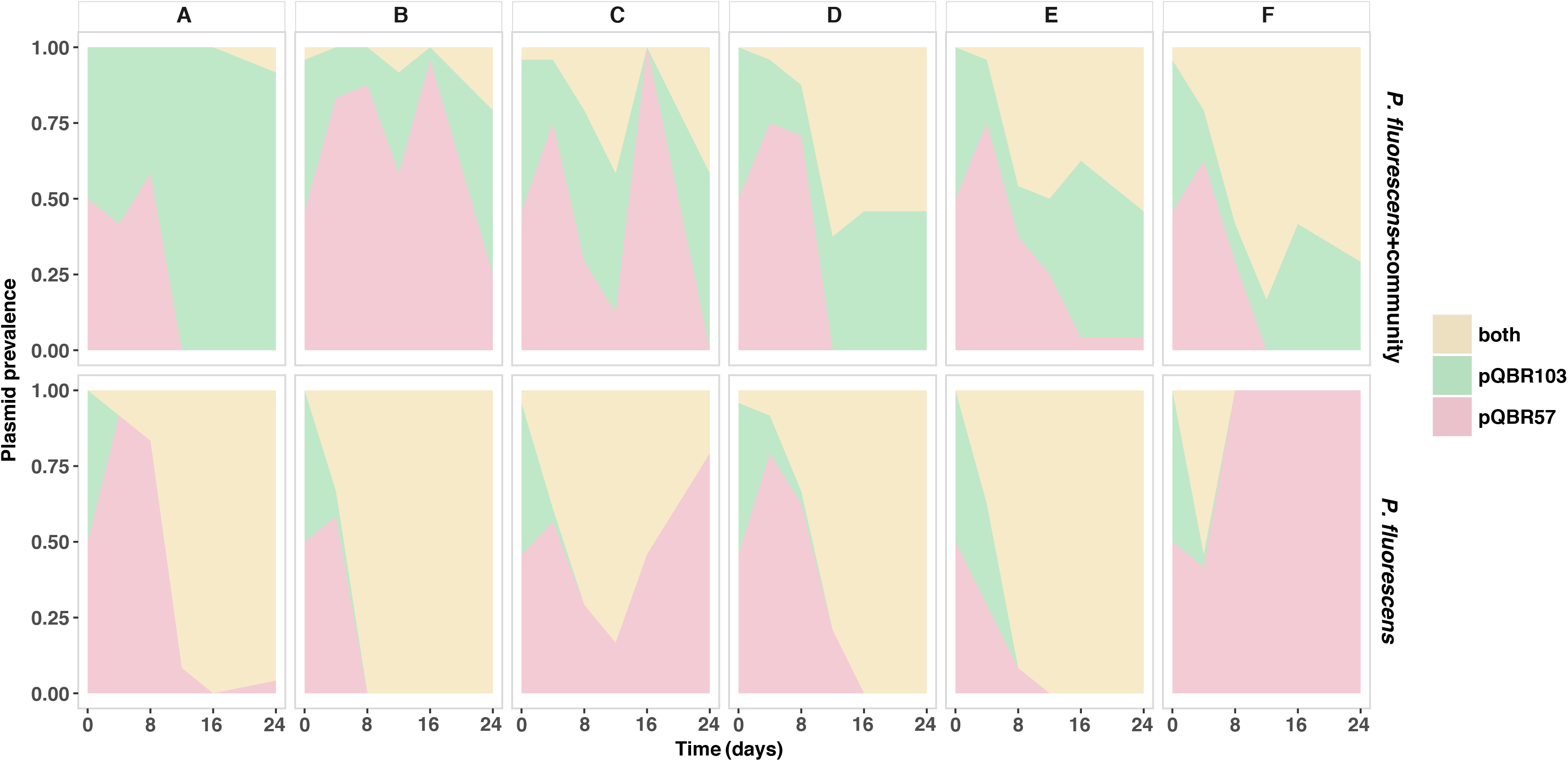
Plasmid ratio and plasmid profile in *P. fluorescens* SBW25. *P. fluorescens +* community panels show the plasmid prevalence in *P. fluorescens* when plasmid-bearing *P. fluorescens* species were co-cultured with the five-species community; *P. fluorescens* panels show the plasmid prevalence in *P. fluorescens* when *P. fluorescens* was cultured as single-species. Co-existence of both, pQBR57 and pQBR103 plasmids (yellow); pQBR103 plasmid (green); pQBR57 plasmid (red).

**Figure 2.**
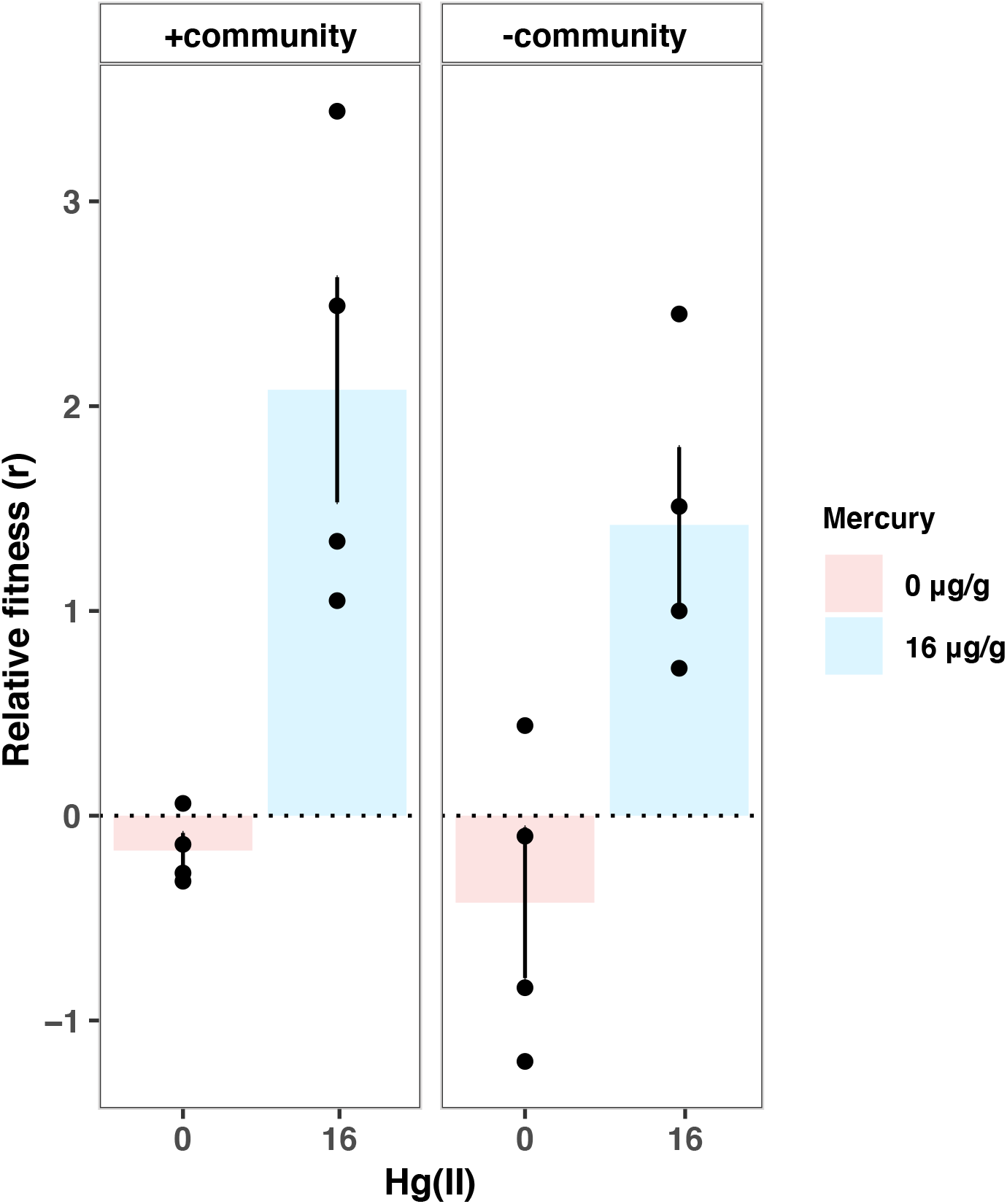
Relative fitness of *P. fluorescens* (pQBR103:pQBR57) in absence and presence of the five-species community. 0 μg g^−1^ Hg(II) (pink), 16 μg g^−1^ Hg(II) (blue). Circles represent the individual data points of four clonal replicates. Error bars represent the SEM of four clonal replicates.

pQBR57 is known to have a far higher conjugation rate than pQBR103 in potting soil [15], therefore it is likely that co-infection would have often resulted from pQBR57 conjugating into cells that already carried pQBR103. This process of infectious transmission through the *P. fluorescens* SBW25 population could have been less efficient in the presence of the background community if, rather than conjugating into *P. fluorescens* SBW25(pQBR103), pQBR57 conjugated into the other *Pseudomonas* species. This is conceptually similar to the dilution effect in epidemiology whereby biodiversity reduces infection risk in a focal species [13]. Consistent with this idea, we observed high levels of mercury resistance in the total community, of which *P. fluorescens* SBW25 made up only ~18% of the total mercury resistant fraction at the end of the experiment, confirming plasmid transmission of the *mer* operon into the other taxa (Figure 3). Within the mercury resistant fraction of the total community, we were able to detect the more highly conjugative plasmid pQBR57, but not pQBR103, at an appreciable frequency (X^2^(2, N=18)=12.176, p=0.002; Figure 4). Together these suggest that, indeed, the transmission of pQBR57 into *P. fluorescens* SBW25(pQBR103) cells was impeded by dilution by the community leading to reduced co-infection of *P. fluorescens* SBW25.

**Figure 3.**
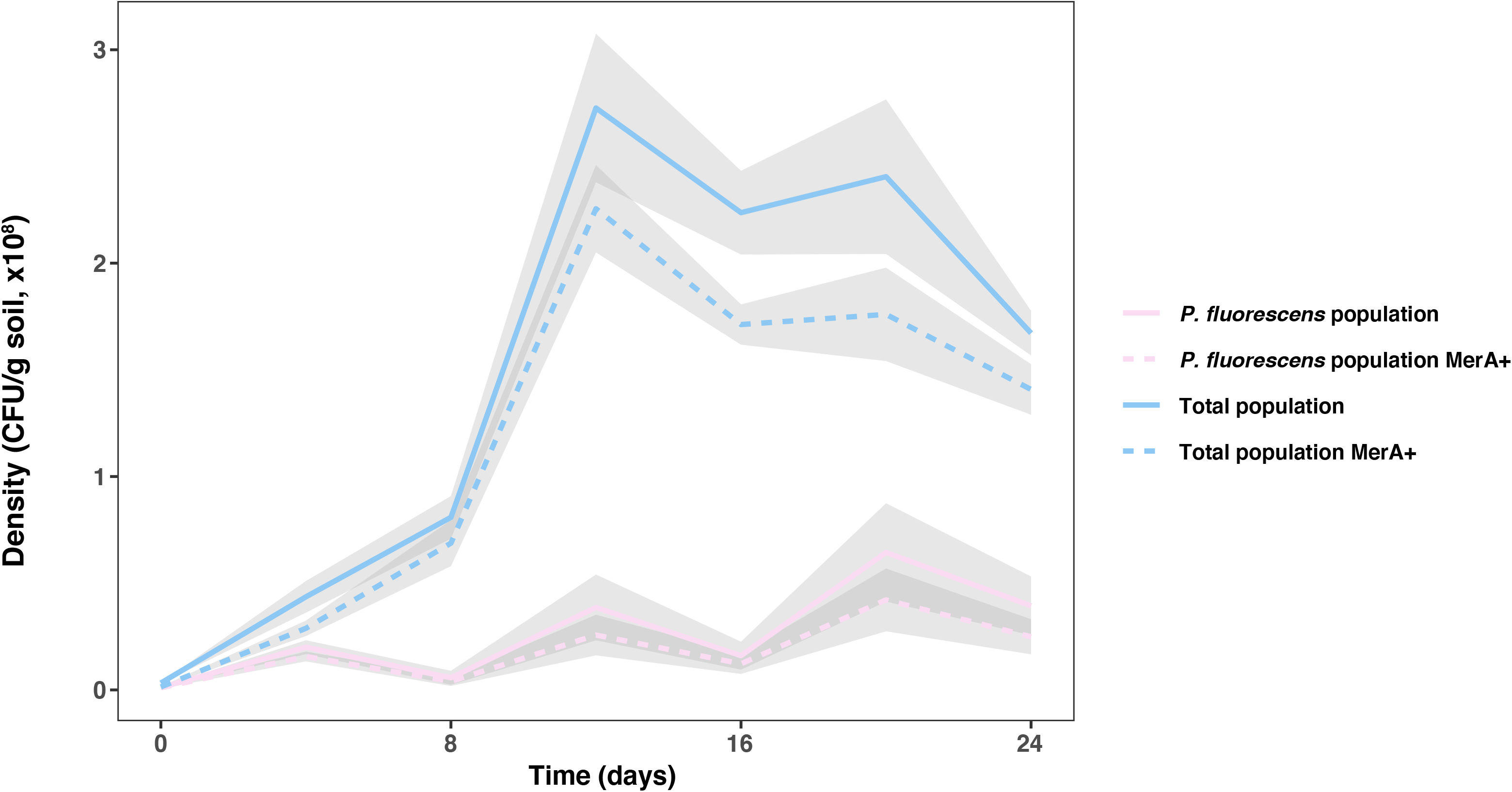
Densities of the total community and of the *P. fluorescens* SBW25 population over time. Solid lines show mean density of the total community (blue) and of the *P. fluorescens* SBW25 population (pink). Dotted lines show mean density of mercury resistant cells in the total community (blue) and the *P. fluorescens* SBW25 population (pink). Grey shaded areas show standard errors (n = 6).

**Figure 4.**
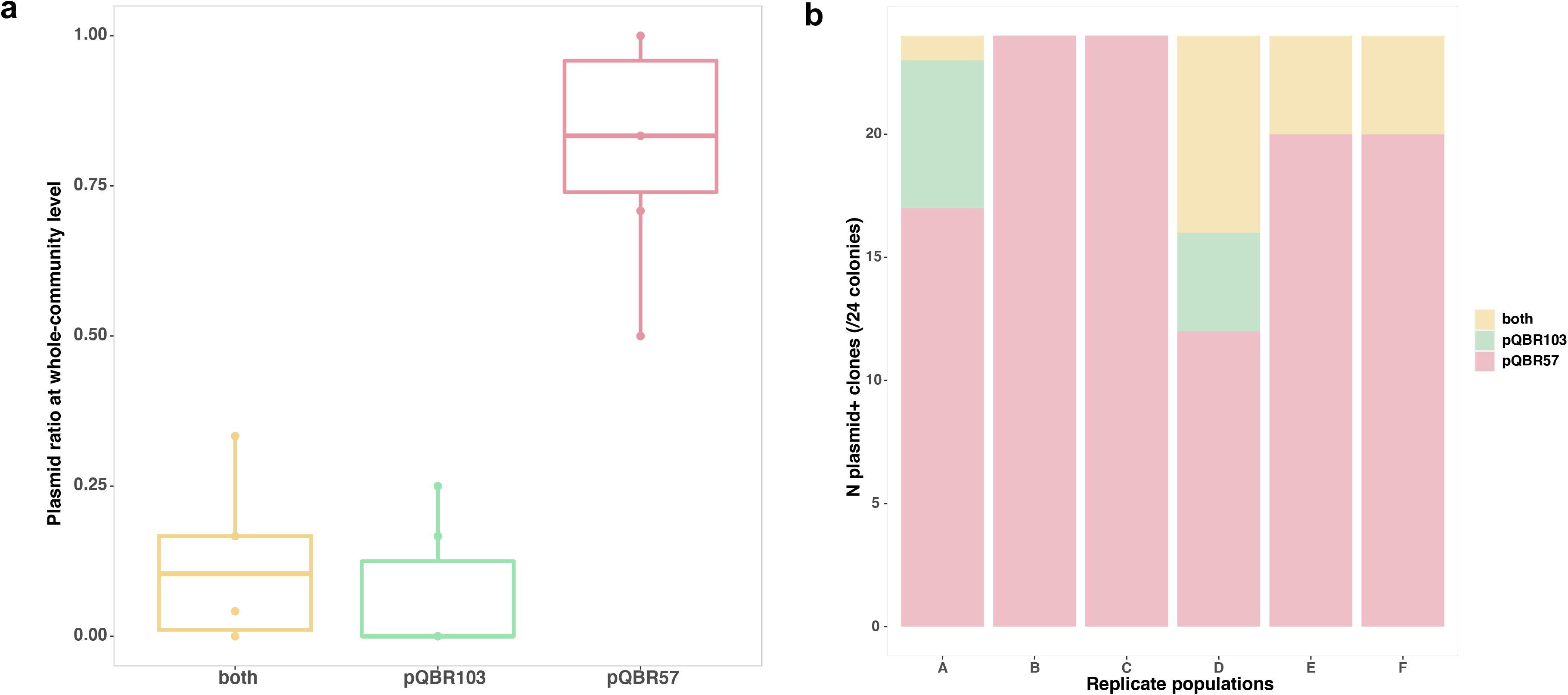
a. Plasmid genotype frequencies in the total community at the end of the experiment. Each box shows the upper and lower quartile, the interquartile range (IDR, length of box) and the median (solid line across the box) of each plasmid genotype frequency in the replicate populations (A-F, n=6). Circles show the outliers of the data. b. Counts of plasmid genotypes in each replicate population (A-F) from twenty-four colonies sampled from the mercury resistant fraction of the total community at the end of the experiment. Co-existence of both, pQBR57 and pQBR103 plasmids (yellow); pQBR103 plasmid (green); pQBR57 plasmid (red).

Finally, we tested whether the rate of conjugation to plasmid-free recipient cells varied depending on whether the donor was singly-infected or co-infected, and whether conjugation rates were affected by resource level to mimic the effects of increased resource competition in more diverse communities. Conjugation rates from all backgrounds — *P. fluorescens* SBW25(pQBR103), *P. fluorescens* SBW25(pQBR57), and *P. fluorescens* SBW25 (pQBR103:pQBR57) — were reduced in diluted media (effect of resource media, X^2^(2, N=22)=16.85, p<0.001; Figure 5; conjugation of pQBR103 was not detectable in medium and low resource media). Consistent with previous studies, conjugation rates from pQBR57-containing backgrounds were far higher than those from *P. fluorescens* SBW25(pQBR103) (Welch’s t-test, t_5.581_ = −14.973, p<0.001), but co-infected donors had a reduced conjugation rate compared to *P. fluorescens* SBW25(pQBR57) donors (Welch’s t-test, t_5.773_= −5.751, p=0.001; Figure 5). These results suggest that co-infection itself may have reduced the rate at which pQBR57 spread in the *P. fluorescens* SBW25 population, and that greater resource competition in the presence of the background community may have reduced the rate of infectious spread of both plasmids.

**Figure 5.**
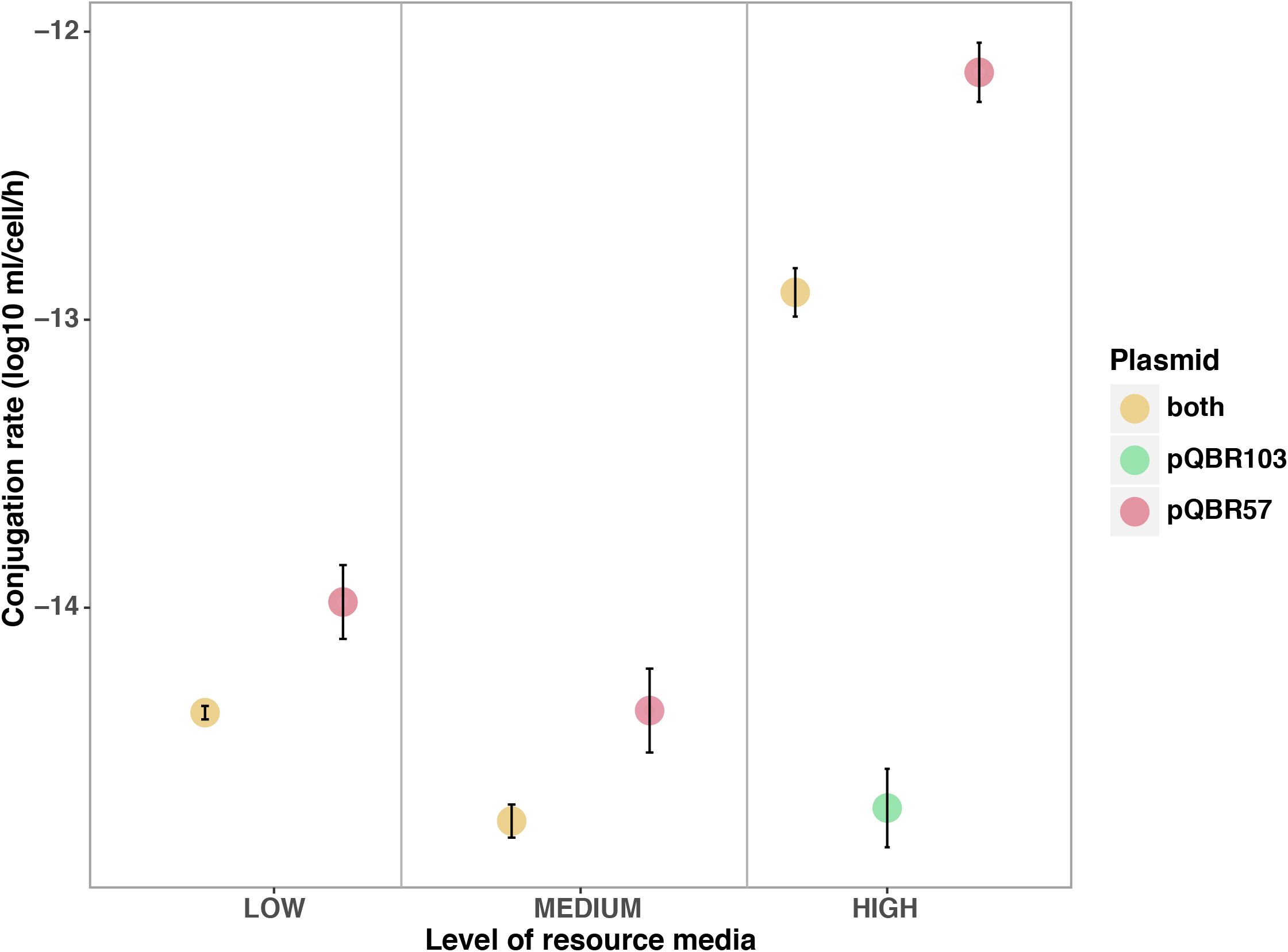
Conjugation rate of *P. fluorescens* (pQBR103:pQBR57), *P. fluorescens* (pQBR103) and *P. fluorescens* (pQBR57) in high, medium and low resource media. Error bars represent the SEM of four clonal replicates.

## Discussion

Using simple soil bacterial communities, we show that plasmid co-infection in a focal host species was reduced in the presence of a community of other bacterial species. This was not caused by differential fitness effects of plasmid-carriage in monocultures versus communities, but rather appears to have been determined by the effect of bacterial species richness on the epidemiology of horizontal transmission of plasmids in the focal host population. Whereas, in monocultures, the highly conjugative plasmid pQBR57 spread into the *P. fluorescens* SBW25(pQBR103) sub-population, in communities this spread was impeded. Detection of pQBR57 at appreciable frequencies in the total community suggests that this effect was due to a substantial fraction of conjugation events leading to the infection of non-SBW25 cells by pQBR57. Because the conjugation rate of pQBR57 may also be lower from other *Pseudomonas* species (e.g., this is known to be the case for *P. putida* [8]), this interspecific conjugation is likely to have had the effect of reducing the overall conjugation rate to *P. fluorescens* SBW25(pQBR103) cells and thus lowering the probability of plasmid co-infection.

Similar to plasmids, the transmission of parasites has often been found to be lower in species-rich communities where a focal species is diluted in the diverse community and therefore has a reduced risk of infection [14, 30–32]. The dilution effect is supported by experimental studies and epidemiological models which suggest that introducing communities of alternative hosts could help to control the transmission of vector-borne diseases caused by parasites (zooprophylaxis) [33–36]. The identity of the introduced host species has important implications in preventing the parasites’ transmission, as different host species are likely to vary in their susceptibility to hosting the parasite [37]. Highly susceptible host species could amplify the disease reservoir of a parasite instead of suppressing it, therefore in order to prevent the dissemination of a parasite, the enrichment of these host species should be restricted in the community [37]. Similar dynamics could apply to plasmids, where host species are known to vary widely in their proficiency to host and transmit plasmids [8].

Parasite epidemiological models also suggest that the species richness of the parasite community can affect the transmission of a focal parasite [38–39]. Both parasite diversity and co-infection have been found to reduce the transmission rate of parasites in a community [38]. Similarly, here we found that the conjugation rate from the donor *P. fluorescens* SBW25(pQBR103:pQBR57) was lower compared with the *P. fluorescens* SBW25(pQBR57) donor. This suggests that plasmid co-infection itself could limit the transmission rate of highly conjugative plasmids, like pQBR57. We speculate that plasmid co-infection affected the plasmid transmission as a result of plasmid-plasmid interactions in the host cell [40]. Co-existing plasmids could trigger a stronger cellular response in the host cell, while the increase in genetic sequence and encoded genes is likely to amplify the physiological and metabolic cost to the host cell, moreover co-infecting plasmids are likely to compete for limited cellular resources (i.e host’s replication factors; [41]). Indeed, we predict that intracellular competition is likely to be more intense between related plasmids, since these will have the greatest overlap in their resource requirements e.g. similar suites of tRNAs.

In nature, bacteria inhabit species-rich communities wherein they co-exist with multiple diverse plasmids [42–43]. The experiments reported here highlight that plasmid dynamics can be affected by both bacterial and plasmid diversity. Plasmids are currently of clinical concern as they often carry and disseminate antimicrobial resistance genes (ARGs) [44]. ARGs are found in bacterial communities colonizing diverse environments where microbial communities can act as resistance reservoirs [45–46]. Expansion of the resistance reservoirs via HGT between bacterial communities is currently an increasing concern [47]. Understanding the transmission dynamics of ARG-encoding plasmids at the community-level is therefore imperative in order to constrain the emergence of resistance in natural microbial communities. This work suggests that plasmid dissemination along with the resistance genes they encode in a focal taxon (e.g. a pathogen) could be limited in more species-rich communities, where plasmid transmission is constrained by the dilution effect.

## Acknowledgements

We would like to thank the late Prof. Stuart Levy for providing the strain *P. fluorescens* Pf0-1 and Dr. Christoph Keel for providing the strain *P. protegens* Pf-5.

